# Multimodal single cell analysis reveals a link between flowering and leaf initiation

**DOI:** 10.64898/2026.09.23.753879

**Authors:** Mark A. A. Minow, Zuhaa Ali, Ankush Sangra, Noah J. Behrendt, Lewis Lukens, Joseph Colasanti, Robert J. Schmitz

## Abstract

*Zea mays* (maize) flowering time is genetically determined and a critical yield determinant. Yet mechanistic understanding of maize flowering remains poor. *Indeterminate1* (*Id1*), a zinc-finger transcription factor (TF), is a monocot-conserved master regulator of maize flowering. Epistasis between *Id1* and the *ZeaCentroradialis*-*Delayed Flowering1* (*Zcn*-*Dlf1*) inductive pathway partly explains ID1 floral control; however, the strong mutant *id1^-^* floral delay is not explained by this pathway alone. To better characterize *Id1* actions, we performed single-cell assay for transposase-accessible chromatin and single nucleus RNA sequencing (scATAC-seq and snRNA-seq) comparing *Id1*^+^ and *id1*^-^ developing leaves. These analyses reveal *id1^-^* chromatin remodeling via TEOSINTE BRANCHED1 CYCLOIDEA PROLIFERATING CELL FACTOR (TCP) and APETALA2/ETHYLENE RESPONSEFACTOR (AP2/ERF) transcription factors and provide candidate direct targets that include *AP2/ERF* genes. These candidate direct targets include the family of *β-glucosidase* genes that lose expression in *id1^-^*. Unexpectedly, CRISPR/Cas9 *β-glucosidase* edits produced plants that phenocopied *terminal ear1^-^* (*te1^-^*) mutants. This phenocopy prompted an investigation into the genetic relationship between *id1^-^*, *te1^-^* and flowering. Surprisingly, *id1^-^ te1^-^* plants exhibited a synergistic floral delay, producing ∼90 leaves before inflorescence production. Beyond highlighting hitherto unappreciated *Te1* autonomous flowering roles, this genetic synergy raises the hypothesis that meristem leaf primordia cessation underpins maize flowering.

## Introduction

Timing of flowering is critical for grain crop productivity, with reproduction being energetically costly and environmentally sensitive. The transition to flowering occurs in the shoot apical meristem (SAM), which then stops producing leaf primordia, and begins reproductive growth. As such, the floral transition and coincidental vegetative growth cessation produces a trade-off; reproductive growth not only uses energy, water and minerals but diverts them from photosynthetic tissue production, limiting total growth potential. Therefore, to maximize yields (Kaufmann, 1961), flowering must be optimized to fully utilize the good growing season while leaving enough time for grain maturity under favourable conditions.

The SAM typically integrates environmental (e.g., photoperiod) and endogenous (e.g., size, age) signals to time flowering. However, temperate *Zea mays* (maize) lost its obligate photoperiodic requirement during Native American adaptation of maize for higher latitude cultivation (Matsuoka *et al*., 2002; Hung *et al*., 2012; Yang *et al*., 2013; Huang *et al*., 2017). This makes temperate maize a ‘day-neutral’ plant that controls flowering predominantly through autonomous floral induction (Coles *et al*., 2010), which relies upon sensing age (Singleton *et al*., 1951; Chuck *et al*., 2007; Yang *et al*., 2023; Poethig and Fouracre, 2024) and developmental maturity (Colasanti *et al*., 1998; Colasanti and Coneva, 2009; Mascheretti *et al*., 2015). As such, temperate maize produces a fixed, genetically determined, number of leaves (Irish and Nelson, 1991; Coles *et al*., 2010; dos Santos *et al*., 2023) before the SAM produces the male inflorescence.

Leaves originate from stem cell outgrowths, known as leaf primordia, which form on the SAM periphery. Leaf primordia formation occurs in a reproducible pattern (phyllotaxy) and rate, with the time between leaf primordia initiation known as the plastochron. Although under complex control (Bassiri *et al*., 1992), grass leaf initiation is partially regulated by the RNA-binding gene *Terminal ear1* (*Te1*) (Veit *et al*., 1998; Kawakatsu *et al*., 2006). Maize *Te1* is expressed in a ‘U-like’ pattern in vegetative SAMs that constrains leaf primordia initiation to physio-normative positions and plastochron (Veit *et al*., 1998). During the maize reproductive transition, leaf initiation stops, and the SAM transitions into sequentially producing inflorescence meristems, tassel branch meristems, spikelet pair meristems, spikelet meristems and then determinate floral meristems (Irish and Nelson, 1991; McSteen *et al*., 2000). All these meristems develop stem cell niches expressing *BARREN INFLORESCENCE3* (*ZmBIF3*) and *CLAVATA3/EMBRYO SURROUNDING REGION-RELATED7* (*ZmCLE7*; respective orthologs of *Arabidopsis thaliana WUSCHEL* and *CLAVATA3*) (Rodriguez-Leal *et al*., 2019; Chen *et al*., 2021; Liu *et al*., 2021; Bang *et al*., 2025; Xu *et al*., 2025) that reorganize cellular divisions and orchestrate reproductive organogenesis. Models of floral induction stress the importance of establishing reproductive identity (Turck *et al*., 2008; Maple *et al*., 2024), but not leaf identity suppression.

Mechanistic understanding of maize floral induction lags other models like *Oryza sativa* (Rice) and Arabidopsis, where flowering signals integrate to induce leaf ‘florigenic’ PHOSPHATIDYL ETHANOLAMINE-BINDING PROTEINs (PEBP) proteins like FT (Araki *et al*., 1998; Abe *et al*., 2005; Wigge *et al*., 2005) and HEADINGDATE3a (HD3a) (Kojima *et al*., 2002). These florigen PEBP proteins move from leaves, via the phloem, to the SAM (Corbesier *et al*., 2007; Tamaki *et al*., 2007) where they physically interact with the bZIP TF, FD (Abe *et al*., 2005; Wigge *et al*., 2005) or *Os*FD1/7 (Taoka *et al*., 2011), then binding the *cis*-regulatory elements (CREs) near floral-identity genes, starting reproductive growth. Maize PEBPs, known as *Zea*CENTRORADIALIS (ZCN) proteins (Danilevskaya *et al*., 2008; Lazakis *et al*., 2011; Meng *et al*., 2011; Stephenson *et al*., 2019; Castelletti *et al*., 2020), and a bZIP transcription factor, DELAYED FLOWERING1 (DLF1), (Muszynski *et al*., 2006; Sun *et al*., 2020) recapitulate the *FT*-*FD* or *HD3a*-OsFD1/7 inductive circuits found in Arabidopsis and Rice (Muszynski *et al*., 2006; Meng *et al*., 2011; Stephenson *et al*., 2019; Castelletti *et al*., 2020). This *Zcn-Dlf1* pathway underpins much maize flowering time variation (Castelletti *et al*., 2020) and *zcn^-^* and *dlf^-^* maize mutants exhibit floral delays in temperate maize (Muszynski *et al*., 2006; Meng *et al*., 2011; Liang *et al*., 2019). However, loss of INDETERMINATE1 (ID1) function, a zinc-finger TF that acts on the autonomous flowering pathway, causes an extreme ∼20 additional leaf floral delay (Singleton, 1946; Colasanti *et al*., 1998), much greater than the delay caused by disrupting the *Zcn-Dlf1* pathway. Epistasis between *id1^-^* and *dlf^-^* places the *Zcn-Dlf* pathway downstream of *Id1* (Muszynski *et al*., 2006; Meng *et al*., 2011). Accordingly, *id1*^-^ floral delay is greater than what is seen in *zcn8^-^* or *dlf1^-^* mutants (*zcn8^-^* ∼1-4 leaf delay (Meng *et al*., 2011; Liang *et al*., 2019); *dlf^-^* ∼8 leaf delay (Muszynski *et al*., 2006)), indicating additional regulators must drive autonomous maize flowering. Indeed, comparing teosinte photoperiodic and autonomous temperate maize induction revealed distinct maize autonomous and photoperiodic leaf gene networks acting in parallel (Minow *et al*., 2018), which supports separate leaf triggers for photoperiodic and autonomous floral induction. Therefore, it appears that *id1^-^* represses *Zcn-Dlf1* floral induction as well as another possible undiscovered autonomous flowering pathway.

*Id1* is expressed in immature leaves away from the SAM, meaning that *Id1*-mediated floral induction involves long distance signalling, which is only partly explained through ZCN actions. Leaf expression profiling revealed *id1^-^* plants alter maize *CONSTANS, CONSTANS-LIKE and TOC* (*CCT*), *APETALA2*/*ETHYLENERESPONSEFACTOR* (AP2/ERF) and *MADS* gene expression (Minow *et al*., 2018). Additionally, *id1^-^* alters carbon metabolism, specifically mature leaf starch:sucrose partitioning, Krebs cycle metabolites (Coneva *et al*., 2012), and immature leaf hemicellulose metabolism gene expression (Minow *et al*., 2018). The functional relationship, if any, between maize flowering and carbon metabolism remains unknown, although links between carbon sensing and flowering exist in Arabidopsis (Wahl *et al*., 2013). *In vitro* DNA affinity assays have revealed an ID1 binding motif (Kozaki *et al*., 2004; O’Malley *et al*., 2016; Zhou *et al*., 2026), and ID1 direct targets are emerging (Wang *et al*., 2023; Zhou *et al*., 2026). Nonetheless, mechanistic understanding of maize autonomous flowering remains mysterious. *MiR3SS* is also altered in expression in *id1^-^* (Minow *et al*., 2018) and represents *in vivo* validated ID1 direct targets (specifically, *MiR3SSc* and *MiR3SSj*) (Wang *et al*., 2023), which, due to the strong ties between phosphate and carbon metabolism (Liu *et al*., 2010), may relate to *Id1* influencing carbon metabolism (Coneva *et al*., 2012). Additionally, *Dhurrinase-related β-Glucosidase* (*β-Glu*) genes are co-expressed with *Id1* in developing leaf tissue and are transcriptionally silent in *id1^-^* (Coneva *et al*., 2007; Minow *et al*., 2018), suggesting they are downstream ID1 targets.

Other genes, like Arabidopsis *APETALA1* (*AP1*), a MADS-family TF, or *APETALA2* (*AP2*), an AP2/ERF TF, control both floral organ identity (Bowman *et al*., 1989; Irish and Sussex, 1990) and flowering time (Mandel and Yanofsky, 1995; Chen, 2004; Yant *et al*., 2010) with *AP1* and *AP2*, respectively, promoting and repressing the floral transition. Homologous maize genes, notably *ZmMADSCS* (Liang *et al*., 2019)*, ZmMADS67* (Sun *et al*., 2020; Zhou *et al*., 2026), and *ZmRELATEDTOAP2.7* (*ZmRap2.7*) (Salvi *et al*., 2007; Liang *et al*., 2019; Sun *et al*., 2020) also control the floral transition. However, models place these regulators on the *Zcn-Dlf1* induction pathway (Liang *et al*., 2019; Zhou *et al*., 2026); as outlined above, this is inconsistent with *ZmRAP2.7*, MADS67 and *MADSCS* serving as the unknown components of ID1-mediated autonomous floral induction. Consistent with maize *id1^-^* suppressing *Zcn-Dlf1* induction, *ZmMADS67* is directly downstream of ID1 and only explains a portion of the *id1^-^* floral delay (Zhou *et al*., 2026). Additionally, *CCT* genes, *ZmCCT9* (Huang *et al*., 2017) and *ZmCCT10* (Hung *et al*., 2012; Stephenson *et al*., 2019), have been identified as potent maize photoperiodic floral repressors, but their expression was lost during maize adaptation to high latitude cultivation, positioning them as unlikely players in *Id1* floral control.

*Id1* orthologs control flowering in diverse monocots (De Riseis *et al*., 2023; Liu *et al*., 2023; Kozaki, 2024) including rice (Wu *et al*., 2008). Unlike maize, rice retains photoperiod sensitivity and its *Id1* ortholog, *Rice Indeterminate1* (*RID1*), affects flowering regardless of photoperiod (Wu *et al*., 2008). Similarly, *Brachypodium distachyon BdID1* is needed for photoperiodic and vernalization-induced flowering (Liu *et al*., 2023), further supporting *Id1* as a monocot-conserved flowering master regulator. RID1 native-promoter-driven chromatin immunoprecipitation sequencing (ChIP-seq) (Zhang *et al*., 2022) analysis revealed direct RID1 targets including an *AP2/ERF* floral repressor, *OsERF136*, and *Headingdate1* (*Hd1)*, a *CCT* floral regulator. However, this ChIP-seq analysis found no RID1 binding near *Hd3a* (Zhang *et al*., 2022); rather, ChIP-seq supported indirect *Hd3a* regulation via RID1 modulation of *OsERF136* and *Hd1* (Zhang *et al*., 2022). This RID1 ChIP-seq supported a ‘TTTGTC’ motif (Zhang *et al*., 2022), which is shorter than that identified for Indeterminate Domain (IDD) family TFs via *in vitro* binding assays (Kozaki *et al*., 2004; O’Malley *et al*., 2016; Zhou *et al*., 2026) and over-expression-based ChIP-seq (Völz *et al*., 2019). The degree to which these mechanisms, and *in vivo* binding motif, are conserved in maize remains unclear.

Here, to examine to what extent *Id1* effects are cell-type constrained, we generated single-cell ATAC-seq (scATAC-seq) and single-nucleus RNA sequencing (snRNA-seq) data from *id1-m1* mutant (*id1^-^*) and *Id1-B73* (*Id1*^+^) immature leaves. This cell-type-resolved comparison highlighted *id1^-^* vasculature changes, notably chromatin accessibility changes in the developing sieve element and expression changes in xylem and phloem vasculature-associated parenchyma. Motif bias within these multimodal data suggests that ID1 mainly alters transcription directly, with most chromatin accessibility changes being driven by secondary effects through TEOSINTE BRANCHED1 CYCLOIDEA PROLIFERATING CELL FACTOR (TCP), AP2/ERF and MYB TF families. By finding differentially expressed genes (DEGs) linked to accessible chromatin regions (ACRs) with RID1 TF binding motifs, we predict candidate direct ID1 targets, which again highlighted *β-Glu* genes. CRISPR/Cas9 edited *β-Glu* function resulted in aberrant plants that phenocopy the *te1* mutant (*te1^-^*). This phenocopy, along with *id1*^-^ *te1*^-^ double mutant phenotypes, hint at interplay between *id1^-^* floral delay and *te1^-^* leaf primordia formation regulation. This unexpected relationship may indicate an underappreciated importance of suppressing leaf identity of SAM during the reproductive transition.

## Results

### *Id1* expression is enriched in vasculature and has signs of a stabilizing regulatory loop

Given the mismatch between the small number of transcriptional changes in *id1^-^* developing leaf (Coneva *et al*., 2007; Minow *et al*., 2018) and the large impact of *id1^-^* on flowering time, we hypothesized that *Id1* impacts are concentrated within rare cell types that are obscured in bulk assays. To investigate *Id1* cell-type-resolved impacts, scATAC-seq measured chromatin accessibility in immature leaves (Figure 1A) of a near isogenic B73 line segregating *id1*^-^ and *Id1*^+^, with two biological replicates per genotype. Filtering nuclei based on Tn5 insertion number, fraction of reads in transcription start sites (FRiT), fraction of reads in peaks (FRiP), organellar reads, and doublet scores (Figure S1) left 5,139 high-quality nuclei. Joint *Id1^+^* and *id1^-^* cell clustering identified 14 clusters with none showing genotypic bias in cell number (Figure 1B; Figure S2). We annotated these clusters into twelve cell-types using *Id1^+^* gene-body chromatin accessibility, a proxy for expression (Marand *et al*., 2021; Tu *et al*., 2022), of cell-specific markers and known cell-type gene ontology (GO) relationships (Figure 1C; Figure S3; Table S1). This cell-type annotation matched the known cellular complements of developing leaves. Curiously, although both genotypes cluster similarly, vasculature marker genes (notably *Sucrose Transporter* family genes) showed aberrant enrichments within *id1^-^* tissues (Figure S4). Similarly, snRNA-seq quality control filtering based on unique molecular identifiers (UMI) and unique gene counts (Figure S5) left 53,506 nuclei and joint *Id1^+^* and *id1^-^* cell clustering produced 11 clusters, which contained enrichment for the expected cell-specific markers and processes. Accordingly, we used marker gene expression to annotate 11 snRNA-seq cell-types (Figure S6). To facilitate comparison across these datasets, we merged respective clusters into similar cell-types (e.g. two scATAC-seq phloem-related clusters and two snRNA-seq clusters were respectively merged to facilitate phloem level comparison), leaving cell-type matched multimodal data from *Id1^+^* and *id1^-^* samples.

**Figure 1.**
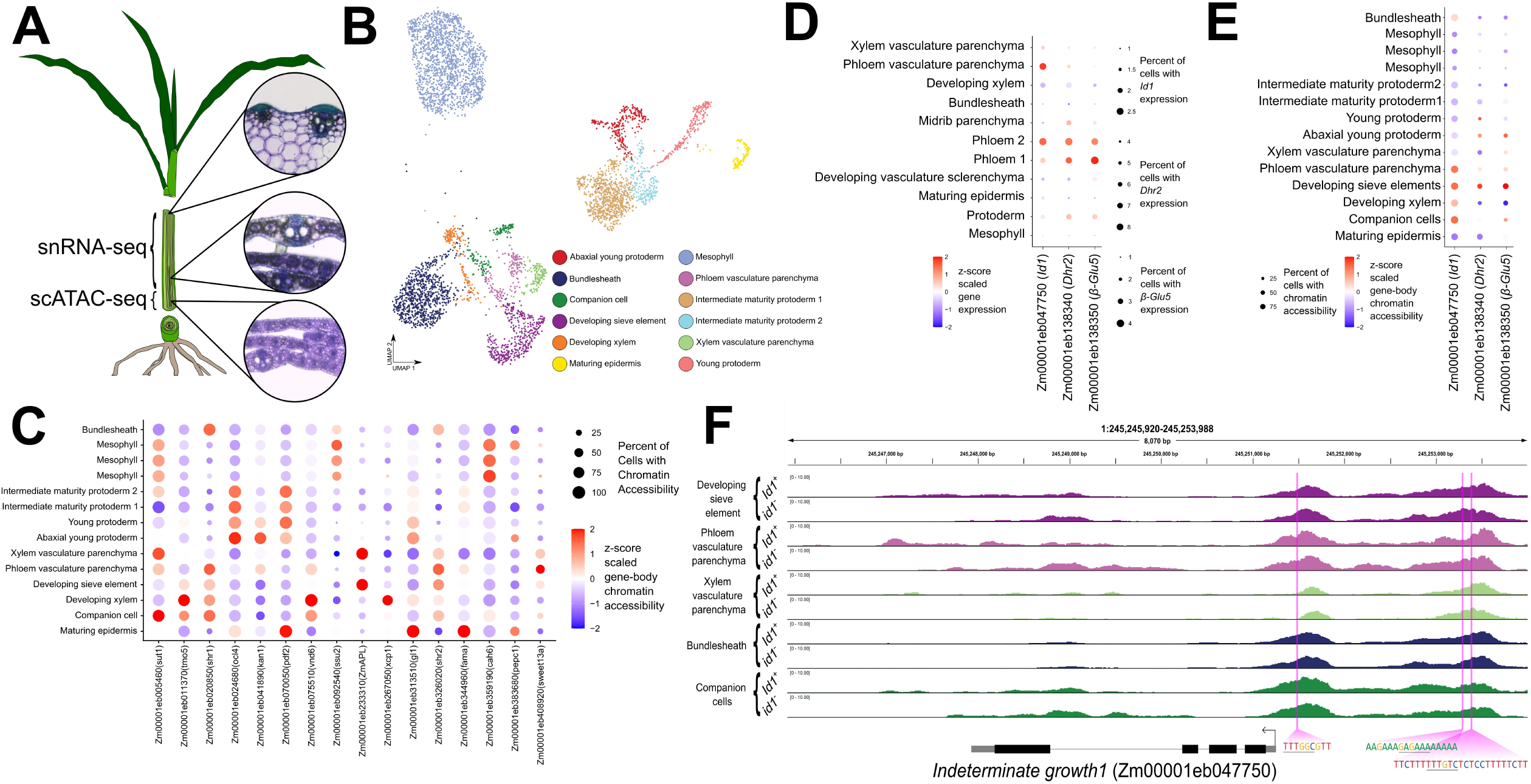
Tandem scATAC and snRNA sequencing sampling of *Id1^+^* and *id1^-^* developing leaves reveals the cellular context of *Id1* expression. (**A**) Two biological replicates of developing leaf whorls were sampled for scATAC-seq and snRNA-seq from B73 near isogenic lines homozygous for either *Id1^+^* (*Id1-B73*) and *id1^-^* (*id1-m1*). To ensure the circadian clock was fully synchronized between the two modalities, the scATAC-seq and snRNA-seq samples were taken contemporaneously from the same individuals; scATAC-seq samples were sampled at a slightly more immature time point than the snRNA-seq samples. The major cellular differences between these two samples were that the midrib parenchyma and vasculature sclerenchyma are developing within the outer leaves in snRNA-seq leaf-whorl and not the scATAC-seq leaf-whorl (circular insets depict this developmental change through representative hand-sections). (**B**) scATAC-seq UMAP dimensionality reductions highlight the cell clusters produced by unsupervised clustering. (**C**) *Id1^+^* z-score scaled gene-body chromatin accessibility, a proxy for transcription, of representative cell-type marker genes that were used to inform the cellular annotation of our cell-clusters. The three mesophyll clusters were merged in subsequent analysis. (**D**) *Id1^+^* z-score scaled cell-type-resolved snRNA-seq gene expression largely corroborate the cellular expression biases inferred from gene-body chromatin accessibility. Note that, despite this bias, *Id1*, *Dhr2* and *β-Glu5* were expressed in all measured immature leaf cell-types. (**E**) *Id1^+^* z-score scaled gene-body chromatin accessibility of *Id1*, and two *β-Glucosidase* genes, *Dhr2* and *β-Glu5* reveal the cellular bias of their native gene-body chromatin accessibility. Again, despite this bias, *Id1*, *Dhr2* and *β-Glu5* had gene-body accessibility in all measured immature leaf cell-types. (**F**) Genome-browser shot showing how *id1^-^* chromatin accessibility was elevated near the *id1-m1* locus, despite no detectable transcript in these samples. Representative cell types are shown for both genotypes. Putative *Id1* binding motif locations, and sequences, are depicted in bright pink (core motif underlined).

Single-cell resolution enabled examination of the cellular context of *Id1* transcription. We employed a ‘pseudobulk’ approach, aggregating reads from individual cells within the same cluster, to generate a cell-type specific profile that resembles a traditional bulk library. *Id1^+^* ‘pseudobulked’ samples exhibited *Id1* nuclear transcripts across all immature leaf cell types (Figure 1D), consistent with past immunohistology. However, expression was particularly high in the phloem and phloem-associated vasculature parenchyma. This was mirrored in the *Id1^+^* scATAC-seq, where gene-body chromatin accessibility was highest in developing vasculature cell-types, except for the xylem-associated parenchyma (Figure 1E). Taken together, this suggests that although widely expressed in leaf cell-types, *Id1* transcription is strongest in the vasculature, particularly the phloem.

These data revealed an increase in *Id1*-linked chromatin accessibility, but not transcripts in *id1^-^* samples (Figure 1F; Table S2). The *id1-m1* mutant allele is caused by a *Ds* insertion (Colasanti *et al*., 1998) that retains a functional promoter and CREs but produces neither functional *Id1* transcripts (Coneva *et al*., 2007; Minow *et al*., 2018; This report) nor protein (Wong and Colasanti, 2007). ACRs upstream of *Id1* contained IDD and RID1 motifs, whereas the *Id1* TSS-proximal ACR contained just the shorter RID1 motif. These motifs, and the *id1^-^* promoter chromatin accessibility increase, hint at a rheostatic transcript feedback loop as seen for other regulatory genes (Bateman, 1998; Rigal *et al*., 2012; Lei *et al*., 2015; Rodriguez-Leal *et al*., 2019; Zhang *et al*., 2024a), where low *Id1* activity is perceived by the cell. In *id1^-^* mutants, this cellular perception appears to elevate *Id1* transcription, however the 1.3kb *Ds* insertion (Colasanti *et al*., 1998) into the third exon prevents the accumulation of functional mRNA, potentially via nonsense mediated decay (Nyikó *et al*., 2013), making this attempted increase in gene expression a futile compensation.

### *id1^-^* altered chromatin accessibility highlights changes near *AP2/ERF* genes in developing leaf phloem

We next found differentially accessible regions (DARs) by comparing *Id1^+^* to *id1^-^* chromatin accessibility. Broadly, cell types exhibited similar chromatin accessibility between *Id1^+^* to *id1^-^* genotypes (Figure 2A). After excluding genes linked to the *id1-m1* introgression (Minow *et al*., 2018, 2021) (Table S3), *Id1^+^* to *id1^-^* comparison revealed between three and 9,455 cell-type DARs (Figure 2B), with predominant changes dispersed across the bundlesheath (9,455 DARs), mesophyll (6,303 DARs), developing sieve elements (5,461 DARs), and one stage of protoderm (intermediate maturity protoderm 1; 4,458 DARs). In these cell types, *id1^-^* coincided with more chromatin accessibility decreases than increases (Figure 2B). Fast gene set enrichment (FGSEA) on DAR-linked genes, revealed very weak gene ontology enrichments centered around changes to primary and secondary metabolism (Figure 2C; Table S4). In aggregate, *id1*^-^-induced chromatin accessibility changes had a marked impact on diverse cell-types, but these changes did not exhibit strong GO enrichments.

**Figure 2.**
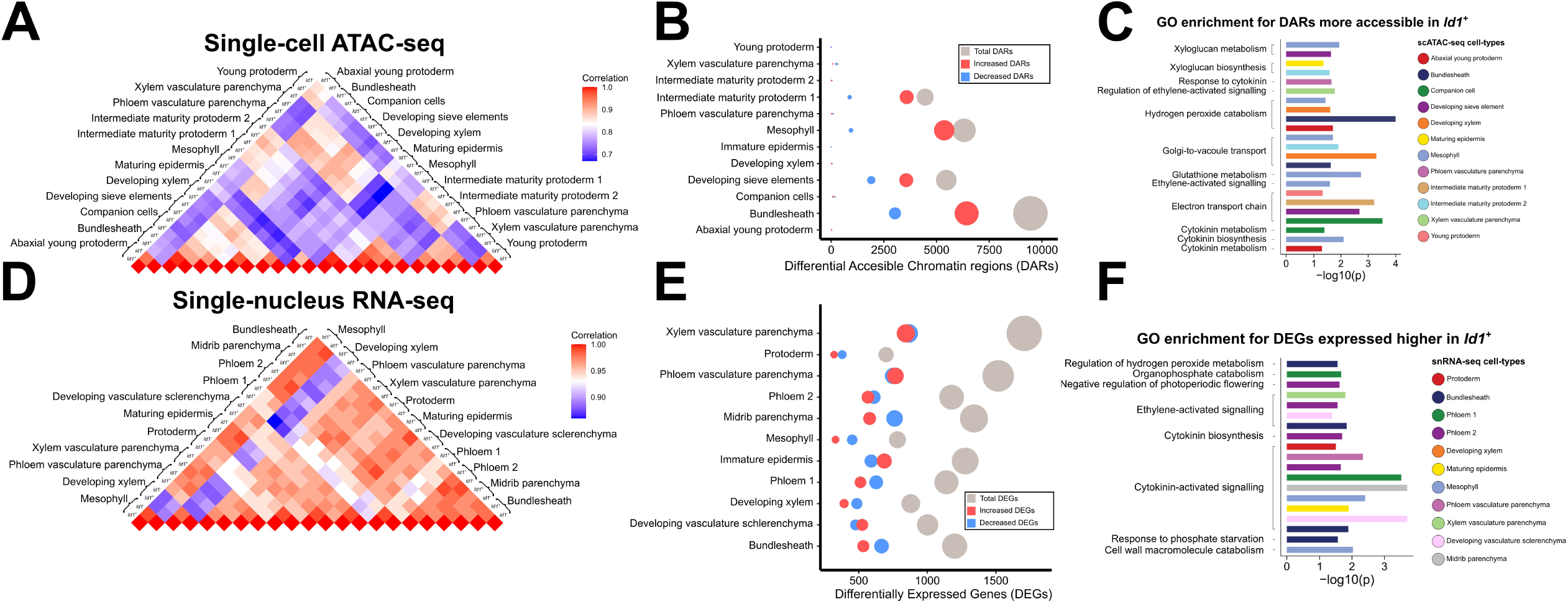
Cell-type-resolved *Id1^+^* to *Id1^-^* comparison highlights a cell-type bias for changes in chromatin accessibility. (**A**) Correlation of cell-type-resolved and ‘pseudobulked’ scATAC-seq indicate that *id1^-^*-induced changes are relatively minor compared to the chromatin accessibility changes induced by cell differentiation. (**B**) Bubble plots illustrating *Id1^+^* to *id1^-^* differentially accessible chromatin regions (DARs) show strong bias towards chromatin accessibility changes in select cellular contexts. Within these changes, more ACRs exhibited higher *Id1^+^* chromatin accessibility (increased DARs), than those higher in the *id1^-^* genotype. (**C**) Gene ontology (GO) enrichment for the genes linked to DARs with higher *Id1^+^* chromatin accessibility supported alterations near hormone and hemicellulose metabolism. (**D**) Correlation of cell-type-resolved, and ‘pseudobulked,’ snRNA-seq likewise support the relatively minor changes in *id1^-^*-induced expression. (**E**) Bubble plots illustrating *Id1^+^* to *id1^-^* differentially expressed genes (DEGs) show that *id1^-^*-induced expression changes are diffuse throughout developing leaf cell-types, with the most pronounced changes in phloem and xylem associated vasculature parenchyma, which contrasts with our DAR analysis. There was no apparent bias towards increased or decreased expression within the *id1^-^*-induced expression changes. (**F**) GO enrichment within DEGs with elevated *Id1^+^* expression highlight alterations to hormones, phosphate metabolism and cell-wall catabolism.

We then examined the specific genes linked to DARs to see the extent to which they could explain known *Id1* impacts on flowering and leaf carbon metabolism. Many DARs were linked to genes associated with xylose-derived hemicellulose metabolism (Table S5). These changes were enriched (binomial test; FDR-adjusted p=7.67e^-06^) in the developing sieve element, where most (binomial test; p=4.36e^-12^) xylose related genes were down regulated in *Id1^+^* tissue compared to *id1^-^*. Some of the largest chromatin accessibility changes were linked to *Glutaredoxin, Thioredoxin* and *Glutathione-S-transferase* genes (Table S5), which influence cellular peptide reductant levels (Holmgren, 1989; Vieira and Rey, 2006; Rouhier *et al*., 2008). These redox-gene-linked ACRs had altered chromatin accessibility in many cell-types, largely matching the cell-type distribution of DARs. Aggregating all cell types, the ACRs linked to redox genes were biased toward lower chromatin accessibility in *Id1^+^* tissue compared to *id1^-^* (binomial test; p=9.32e^-22^). Together, these DARs highlight alterations to hemicellulose metabolism, especially in the developing phloem, and more cell-type agnostic increases of peptide reductants in the *id1^-^* immature leaf.

DARs linked to genes from families that regulate the floral transition were also discovered. These DARs were linked to *SPL*-like, *MADS*-box, and, most commonly, *AP2/ERF*-family genes (98 DARs linked to 81 *AP2/ERF* genes; Table S6). DARs near SPL and MADS genes had weak cell-type enrichments, instead largely reflected cell types that had the most DARs, specifically the bundlesheath, mesophyll, developing sieve elements and one stage of immature protoderm. In contrast, DARs linked to *AP2/ERF* genes were strongly biased towards the developing sieve element (binomial test; FDR-adjusted p=7.47e^-11^), where they were all down regulated in *Id1^+^* compared to *id1^-^* (binomial test; p=5.29e^-23^). This is consistent with phloem *AP2/ERF* floral repression being more active in *id1^-^*. Together, DAR examination suggests ID1, directly or indirectly, influences ACRs near *SPL*, *MADS* and *AP2/ERF* genes, and that developing phloem *AP2/ERF*-linked chromatin accessibility increases in *id1^-^*.

### *id1^-^* alters gene expression throughout the developing leaf, with pronounced vasculature parenchyma impacts

We compared *Id1^+^* and *id1^-^* cell-type gene expression via snRNA-seq to identify DEGs (Table S7). Mirroring the DAR analysis, most cell types were largely similar between *Id1^+^* and *id1^-^* genotypes (Figure 2D). Cell-types with the most pronounced expression changes largely differed from those with the most prevalent chromatin accessibility changes, and there was no apparent genotypic bias towards increased or decreased gene expression (Figure 2E). Cell types with the most DEGs were the xylem vasculature parenchyma (1,707 DEGs) and phloem vasculature parenchyma (1,517 DEGs). Xylem-associated vasculature parenchyma FGSEA revealed minor enrichments for altered ion related processes. FGSEA of *Id1^+^* up-regulated genes (Figure 2F) highlighted cytokinin-activated signalling in the phloem vasculature parenchyma (Fisher’s exact test; p=4.61e^-3^), and all other cell-types except for developing xylem and xylem vasculature parenchyma. These cell-type-resolved transcriptomes emphasize the impacts of loss of *Id1* activity on vasculature parenchyma transcription, despite relatively small effects on chromatin accessibility in these cell-types.

Individual gene expression changes mirrored previous bulk transcriptomics (Coneva *et al*., 2007; Minow *et al*., 2018). For example, several *β-Glucosidase* genes exhibited dramatically reduced expression in *id1^-^* tissuesacross all cell types. Additionally, a newly annotated gene, *Xylose isomerase* (*XylA*), which putatively interconverts D-Xylose to D-Xylulose or D-Glucose to D-Fructose (Stevis and Ho, 1985), had dramatically higher expression in all *Id1^+^* cell-types (Table S7). Related to *XylA*, *Xylulose kinase1* (Zm00001eb094750) is highly expressed in developing leaves (Woodhouse *et al*., 2022), but unchanged in our comparison. Unlike previous bulk transcriptomics (Minow *et al*., 2018), expression changes in *Glycosyl hydrolase16* enzymes, putative hemicellulose remodelers, were mild and limited to mesophyll, phloem, epidermis, xylem-associated vasculature parenchyma and putative midrib parenchyma cell-types. *Glutaredoxin, thioredoxin* and *glutathione-S-transferase* genes, which relate to cellular redox control, exhibited cell-type distinct expression differences. These transcriptional changes hint at *id1*^-^ altered glycoside metabolism with consequences on immature leaf redox pools.

We next interrogated expression profiles for known floral regulator gene families (Table S8). This revealed cell-type-level expression changes for 72 *AP2/ERF* genes, including *ZmAP2-ERF-TF115* (*ZmEreb115*; Zm00001eb318910; GRMZM2G069146), which was previously detected as an *id1^-^* DEG in bulk RNA-seq (Minow *et al*., 2018). However, none of these differentially expressed *AP2/ERF* genes appeared orthologous to the RID1 target *OsERF136* (Zhang *et al*., 2022). MADS floral regulators were also DEGs, including *ZmMADS1* (Zm00001eb403750) in the phloem and *ZmMADS67* (Zm00001eb327040) in phloem, xylem vasculature parenchyma, epidermis, developing sclerenchyma and putative midrib parenchyma cells. Mirroring previous bulk profiling (Minow *et al*., 2018), we observed expression changes in 25 *CCT* genes, further supporting a relationship between *Id1* and the circadian clock. However, expression changes of these putative floral regulators were not cell-type biased (Table S9), suggesting *Id1* control of floral gene expression is spread throughout developing leaf cell types.

### *MiR399c* and *MiR399j* show no signs of *in vivo* ID1 regulation of chromatin accessibility

Since the maize miRNA-encoding genes, *MiR3SSc* and *MiR3SSj* are confirmed direct ID1 targets (Wang *et al*., 2023), we investigated cell-type-resolved *MiR3SS* expression and chromatin accessibility. However, microRNAs (miRNAs) are not well represented in the current reference annotation (B73 NAM V5.0), which omitted these miRNA family members in our global comparisons. Accordingly, we specifically examined cell-type-resolved chromatin accessibility and expression at *MiR3SSc* (chr6:171179057-171182263) and *MiR3SSj* (chr9:26295469-26298643). At these loci expression was low and beneath the detection limit of the snRNA-seq (data not shown), potentially reflecting rapid processing of pre-miRNA transcripts into miRNA duplexes (Yu *et al*., 2026) or low expression in the sampled tissues. Gene body chromatin accessibility over *MiR3SSc* and *MiR3SSj* was also low, but detectable in the scATAC-seq, with *MiR3SSj* exhibiting perceptible chromatin accessibility in many cell types (Figure 3A). Consistent with a role in vasculature-based phosphate control (Pant *et al*., 2008), *MiR3SS* isoform chromatin accessibility was prevalent within vasculature cell types (Figure 3A), which actively manage systemic solutes, like phosphate (Lin *et al*., 2014; Zhang *et al*., 2014). However, unlike *MiR3SS* levels (Minow *et al*., 2018; Wang *et al*., 2023), no differences were seen in *MiR3SSc* and *MiR3SSj* gene body chromatin accessibility between *Id1* and *id1^-^* genotypes (Figure 3B), suggesting minimal effects of *id1^-^* loss of function on chromatin structure at these loci. Since *MiR3SSj* exhibited higher chromatin accessibility, we examined it more closely. Here, only one of the two *MiR3SSj-*linked ID1 binding sites assayed in Wang et al. (2023) was within accessible chromatin (Figure 3C), suggesting this proximal ID1 binding site may have greater *in vivo* importance than the inaccessible motif. Together, our data indicate that, at least for these direct-target miRNAs, that ID1 can modulate expression without significantly altering chromatin accessibility.

**Figure 3.**
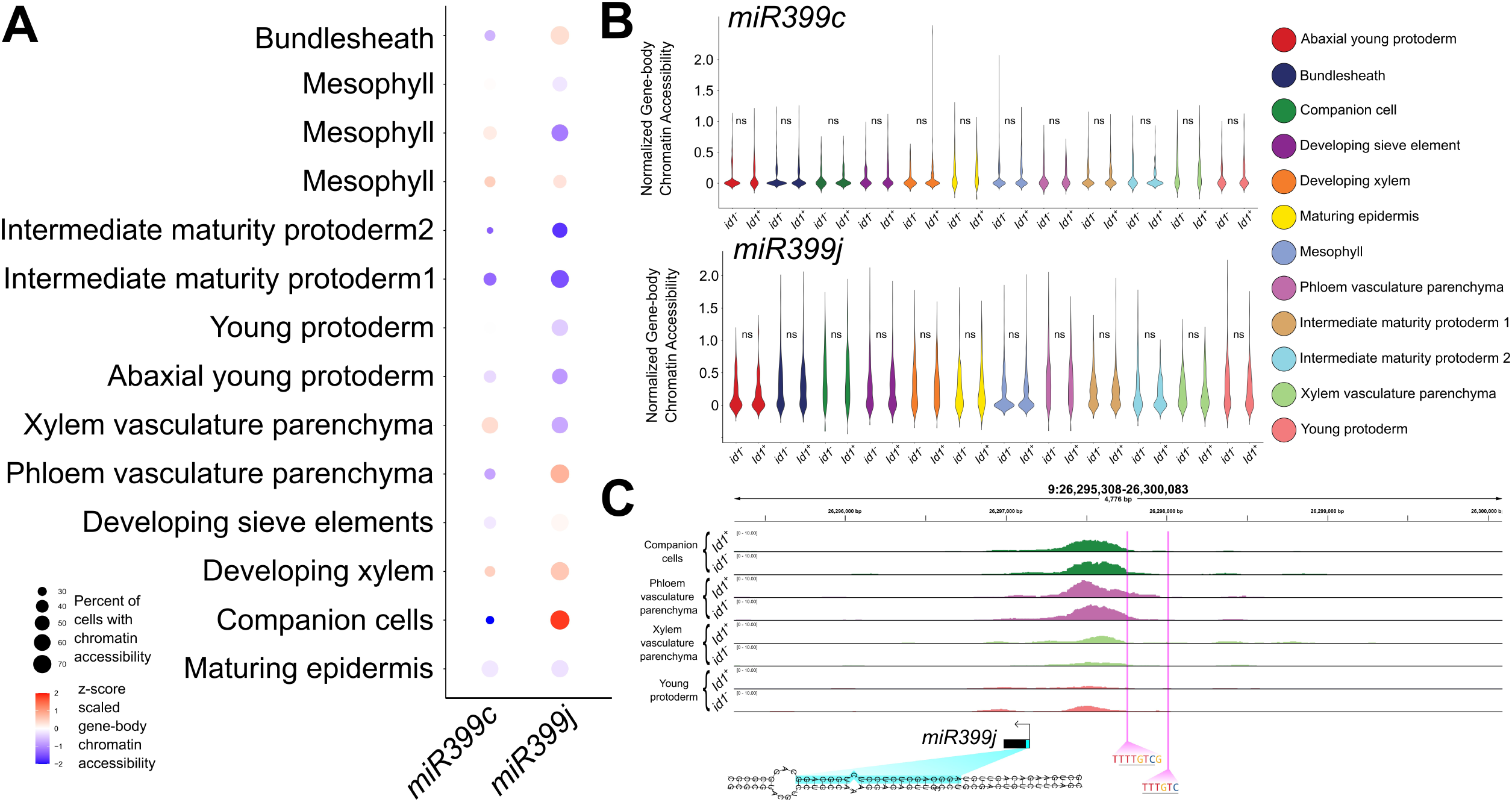
Chromatin accessibility linked to *Mir333c* and *Mir333j* was unaltered by loss of *Id1* function. (**A**) Gene-body chromatin accessibility of *MiR3SSc* and *MiR3SSj* were both biased towards vasculature cell-types, although the extent of that bias varied by miRNA isoform. (**B**) Depiction of miRNA-linked chromatin accessibility for *MiR3SSc* (top) and *MiR3SSj* (bottom) for both *Id1^+^* and *id1^-^* cell-types. Pairwise comparisons between genotypes supported no significant alteration to *MiR3SS* chromatin accessibility (ns; p>0.05; t test), for either isoform. (**C**) Genome browser-shot examination likewise supported similar accessibility at *MiR3SSj*. Representative cell-types shown. Only one of the two previously assayed ID1 binding sites (sequences and locations depicted in bright pink; core motif underlined) was within accessible chromatin in the developing leaf. The *MiR3SSj* locus was inferred from a Maize V4 genome lift-over, and confirmed via RNA-fold, which supported a proper hairpin over the active miRNA site (highlighted in cyan). *MiR3SSj* was selected for this depiction, as it was more consistently in accessible chromatin in our sample.

### Loss of *Id1* function alters TCP, ERF/AP2 and MYB motif chromatin accessibility, while the RID1 motif is enriched near transcriptional changes

Since the ID1 binding motif is known (Kozaki *et al*., 2004; O’Malley *et al*., 2016; Zhang *et al*., 2022), we investigated IDD/RID1 motif chromatin accessibility in both *Id1^+^* and *id1^-^* contexts (Data S1). Across all cell types, IDD/RID1 motif chromatin accessibility was strongly enriched in the mesophyll, with weaker enrichment seen in vasculature cells (Figure 4A; Figure S7), as reported in maize and rice (Marand *et al*., 2021; Yan *et al*., 2025b). Next, we examined motif enrichment in *id1^-^*-induced chromatin accessibility changes (Figure 4B; Figure S8). Surprisingly, IDD/RID1 motifs were not the predominant signal in genotypic chromatin accessibility differences. Further, comparing the proportion of DARs with an IDD/RID1 motif to that predicted by chance, revealed IDD/RID1 motif depletion in the DARs (Table S10; Figure 4C). Rather, TCP and AP2/ERF TF motifs had the largest changes in chromatin accessibility between *id1^-^* and *Id1^+^* cells (Figure 4B; Figure S8). TCP motifs were mostly enriched within ACRs with heightened *id1^-^* chromatin accessibility. In contrast, AP2/ERF TF motif chromatin accessibility changes were more evenly split between higher or lower *id1^-^* chromatin accessibility depending on cell type. MYB TF family motifs also exhibited altered chromatin accessibility, predominantly in ACRs with decreased *id1^-^* chromatin accessibility. Together, the lack of IDD/RID1 motifs within *id1^-^* chromatin accessibility changes indicate that *id1^-^* predominantly alters chromatin accessibility through indirect effects, potentially by altering TCP, AP2/ERF and MYB TF activities.

**Figure 4.**
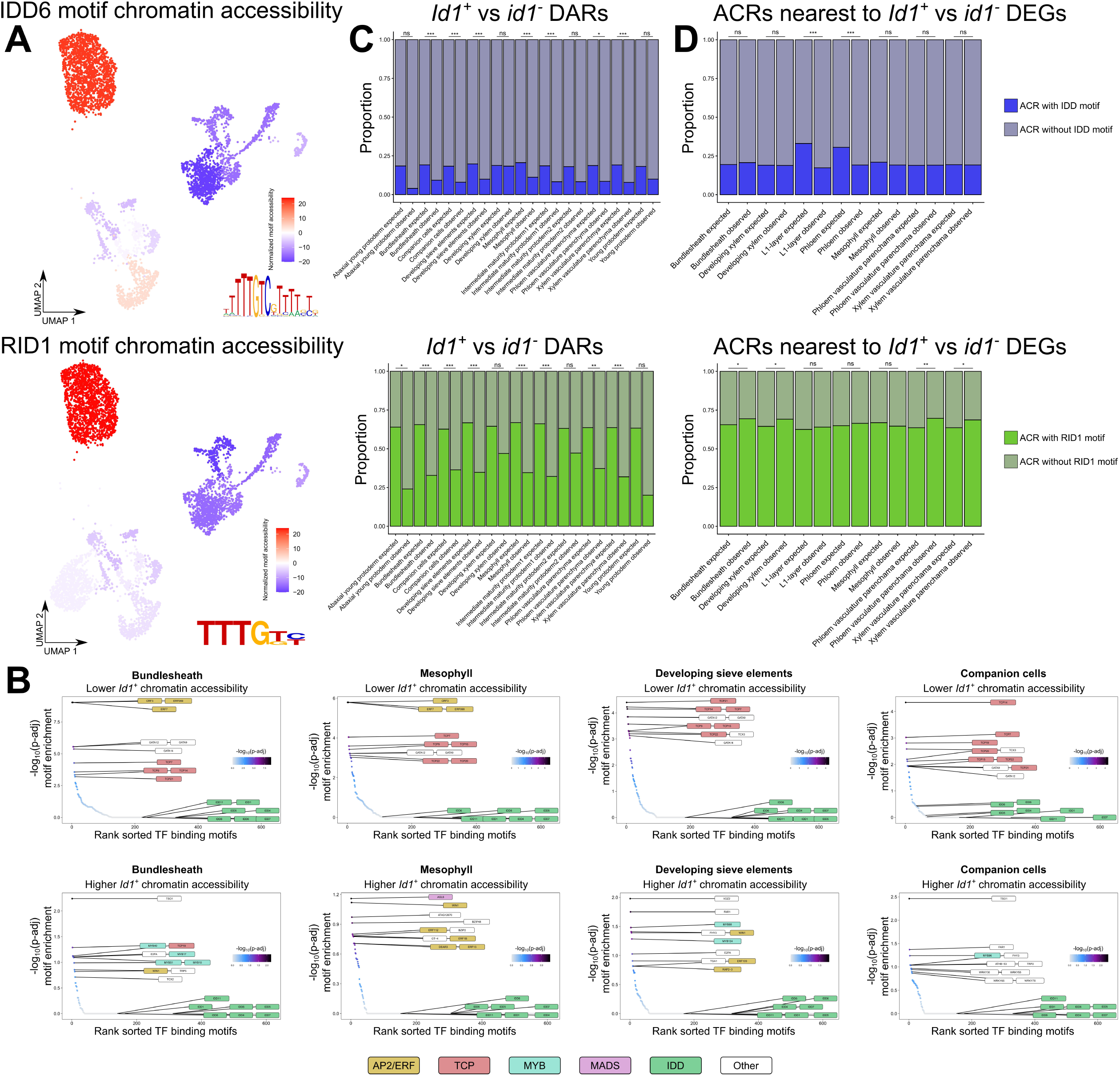
*Id1^+^* to *Id1^-^* changes in TF binding motif chromatin accessibility were centered around AP2/ERF, TCP and MYB TF binding motifs and not IDD binding motifs. (A) Global comparison of cell-type bulked IDD6 (JASPAR motif ID: MA2348.1) and RID1 TF binding motif chromatin accessibility highlight strong enrichment with the mesophyll, with a lesser enrichment within bundlesheath and vasculature cell-types. IDD6 was chosen as a representative of the established IDD-family motifs, as the family largely behaved similarly. Sequence logos of the two motifs are shown in the bottom right corner of each UMAP. (B) ChromVAR TF binding motif enrichments showing which motifs had lower and higher *Id1^+^* chromatin accessibility (top and bottom row, respectively), for four representative cell types. AP2/ERF, TCP and MYB TF family motifs were most well represented in these *Id1^+^* to *id1^-^* chromatin accessibility changes. In contrast, all JASPAR-annotated IDD motifs were poorly represented in these changes. Note IDD6 appears twice, as two similar, but distinct, motifs exist for this TF in the JASPAR database. (C) Comparing the proportion of DARs with an IDD or RID1 motif (top and bottom, respectively) to that predicted due to chance (Expected vs Observed). This comparison supported consistent depletion of IDD and RID1-containing ACRs in the observed DARs. This further supports the notion that the observed chromatin accessibility changes are not being directed by binding to either the RID1 or IDD motif. (**D**) Comparing the proportion of DEGs with a linked ACR containing either IDD or RID1 motif (top and bottom, respectively) to that predicted due to chance (Expected vs Observed). This comparison did reveal a mild enrichment in RID1 motif, but not IDD motif, presence within ACRs near DEGs from select cell-types. *** p<0.001; ** p<0.01, * p<0.05, ns p>0.05; t test. Please refer to Figure 1 for the cell-type identities on the scATAC-seq UMAP.

Investigating how TCP and AP2/ERF TFs might contribute to *id1^-^* perturbed chromatin accessibility, we looked at the genes linked to DARs containing either motif. Specifically, we looked at the FGSEA of genes linked to all DARs containing TCP or AP2/ERF motifs (Table S11). Aggregating cell-types, DARs with TCP motifs were linked to TFs (GO:0003700,DNA-binding TF activity; Fisher’s Exact test; p=7.80e-11) while DARs with AP2 motifs exhibited weaker enrichments (Fisher’s Exact test; p>=1.3e^-3^) largely centered around changes to cellular energy/carbon metabolism. This suggests that *id1^-^* potentially partitions TF and energy metabolism changes via TCP and AP2/ERF TF families, respectively.

We next examined the motifs in ACRs near DEGs to see if the transcriptional consequences of *id1^-^* were more enriched for nearby IDD/RID1 motifs. To do this, we again compared the proportion of DEG-linked ACRs with IDD/RID1 motifs to that predicted by chance. Unlike the DAR analysis, this revealed mild motif enrichment in DEG-linked ACRs, but just for the RID1 motif (binomial test; p ∼0.01-3.0e^-4^; Figure 4D; Table S12), and only in bundlesheath, developing xylem, xylem vasculature parenchyma, and phloem vasculature parenchyma cells. Taken together, direct ID1 transcriptional regulation is most supported by our data, although secondary effects undoubtedly mask direct ID1 effects on both gene expression and chromatin accessibility.

### Candidate ID1 direct targets encompass both floral and carbohydrate-associated genes

Maize direct ID1 targets, and how they orchestrate flowering, remain poorly understood. Our data identify candidate direct targets by revealing which *id1^-^* DEGs have linked ACRs with IDD/RID1 motifs. Given the better RID1 DEG-linked ACR motif enrichment, we focused on DEGs linked to RID1-containing ACRs. We also required a TTTGKC motif (K represents a G or T), as *in vitro* binding was largely lost without this 3’ cytosine (Kozaki *et al*., 2004). Accordingly, we identified 7,459 unique candidate ID1 target genes (Table S13). Candidate direct targets included MADS67 (Zm00001eb327040), 24 *CCT* genes, and *ZCN11* (Zm00001eb281410). Encouragingly, *MADS67* has molecular evidence supporting direct ID1 binding (Zhou *et al*., 2026). Additionally, 68 *AP2-ERF* TFs were differentially expressed and were linked to RID1-containing ACRs, consistent with the DAR motif enrichments and RID1 ChIP-seq (Zhang *et al*., 2022). These candidate direct targets were mildly biased towards increased expression in *Id1^+^* vs *id1^-^* compared to chance (∼52.4% upregulated; binomial test; p=7.67e^-8^). However, given the validated maize direct ID1 targets are both activated (MADS67; SBP20) (Zhou *et al*., 2026) and repressed by ID1 (*MiR3SSc* and *MiR3SSj*) (Wang *et al*., 2023), it is possible ID1 impacts nearby transcription bidirectionally, potentially through different TF binding partners (Wang *et al*., 2023; Zhou *et al*., 2026). Indeed, these candidate *AP2/ERF* targets had inconsistent *Id1^+^* vs *id1^-^* expression changes, suggesting complex locus-specific regulation, likely conflating both direct ID1 effects and indirect consequences.

Other ID1 candidate direct targets included genes associated with carbon metabolism (Table S13). This included *XylA*, a cell wall invertase (*miniature seed1*, Zm00001eb083790) and a starch-debranching enzyme (*sugary1*, Zm00001eb174590). Additionally, all glucosidase genes known to lose expression in *id1^-^* (Zm00001eb138340, *dhurrinase2*, *Dhr2*; Zm00001eb138350, *β-Glucosidase5*, *β-Glu5*; Zm00001eb411430, *β-Glucosidase3*, *β-Glu3*) had linked ACRs with RID1 motifs, including a distal ACR linked to *Dhr2* and *β-Glu5* (Figure 5A). This provides a plausible direct regulatory mechanism for *id1^-^* to control *β-Glu* expression, although the roles of these *β-Glu* genes in metabolism and flowering remain uncharacterized.

**Figure 5.**
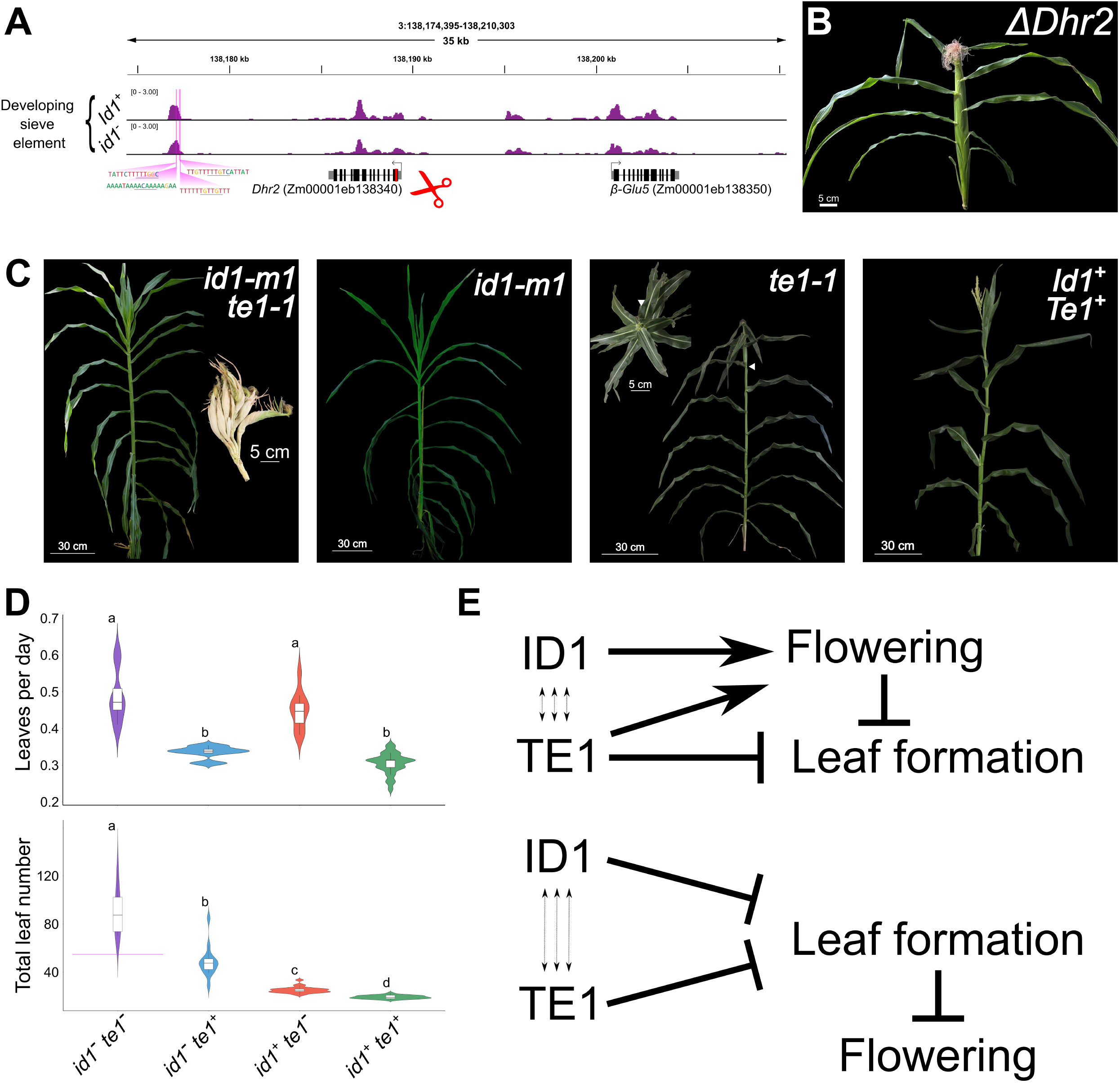
*Dhr2* in-frame CRISPR/CasG editing and double mutant analysis unveil *Id1* and *Te1* synergistically control autonomous flowering. (A) Genome browser-shot of developing sieve element chromatin accessibility near *Dhr2* and *β-Glu5* which are duplicated genes present in an inverted repeat. These two genes lose all expression in *id1^-^* backgrounds and contain several putative ID1 binding sites in a distal ACR (sequences and locations depicted in bright pink; core motif underlined). Red bar and scissors depict the location of repeated CRISPR/Cas9 edits, which produced in-frame *Dhr2* edits (*ΔDhr2;* 66-bp in-frame deletion starting at the 70th coding nucleotide) five independent times. Developing sieve elements were chosen for depiction, as this is the cell-type with the highest expression for these genes. (B) All five independent *ΔDhr2* edits produced a dominant plant phenotype characterized by extreme dwarfism, weak vigour, and production of a fully feminized terminal ear enclosed in husk leaves. *ΔDhr2* plants phenocopied *te1^-^* phenotypes, although with reduced vigour and more penetrative tassel feminization. Note the extreme dwarfism that allows the entire plant to be depicted in B. (**C**) F2 whole plant morphology produced from either *id1^-^ te1^-^* double mutants, either single mutant, or *Id1^+^ Te1^+^* plants. Morphologically, *id1^-^ te1^-^* double mutants resembled that of *id1^-^* single mutants throughout vegetative development. However, the male inflorescence was replaced with a highly branched terminal ear structure (left-most inset). As is typical of *id1^-^* single mutants, no ears were produced in any *id1^-^* background. *te1^-^* single mutants displayed their eponymous terminal ear phenotype, although some masculinized flowers were observed in their apical inflorescence. *te1^-^* plants transitioned to a spiral phyllotaxy at ∼leaf 18 (denoted by the white arrowhead; same leaf is highlighted in both the profile and middle-right inset overhead image). (**D**) F2 violin box plots illustrating the leaves produced per day (a measure of plastochron) and total leaf number (a measure of developmental flowering time) for *id1^-^ te1^-^* double mutants, both single mutants, and *Id1^+^ Te1^+^* plants. Leaf production rate was entirely controlled by *Te1* status, with *te1^-^* single mutant resembling the *id1^-^ te1^-^* double mutant and *id1^-^* single mutants resembling *Id1^+^ Te1^+^* plants. However, total leaf number demonstrated a clear synergistic delay in *id1^-^ te1^-^* plants, which produced ∼90 leaves before flowering, which far exceeded that predicted by additive effects (bright pink bar in *id1^-^ te1^-^* genotype). Letters denote statistically identical groups (p>0.05; one-way ANOVA). (**E**) Two genetic models that explain the observed *id1^-^ te1^-^* synergistic floral delay. (Top) *Id1* and *Te1* could both influence flowering, with *Te1* controlling leaf initiation independently. (Bottom) Alternatively, *Te1* and *Id1* may both repress leaf formation, with this repression required for the maize floral transition. In either case, reciprocal pathway crosstalk (triple dashed arrows) is thought to allow pathway compensation in either *te1^-^* or *id1^-^* single mutant backgrounds.

### CRISPR/CasG editing of *Dhurrinase2* phenocopies *terminal ear1* mutation

Past and present expression profiling analyses highlight the relationship between *Id1* and a family of *β-Glu* genes (Coneva *et al*., 2007; Minow *et al*., 2018). Examining cellular *Dhr2* and *β-Glu5* expression, supported near ubiquitous expression in immature leaf cells, with the highest gene body chromatin accessibility and expression in the developing sieve element (Figure 1D). This phloem enrichment is intriguing, as it is the tissue that links the developing leaf to the SAM, putatively transmitting *Id1* floral inductive signals.

*Dhr2* and *β-Glu5* are highly similar (>91% amino acid identity) and tightly linked in an inverted repeat (chr3:138185404-138204545). This presumed functional redundancy and tight linkage precluded functional characterization through reverse genetics using classical mutagens. However, CRISPR/Cas9 editing can genetically couple *Dhr2* and *β-Glu5* mutations onto one chromosome. To this end, we began generating both single and double *Dhr2* and *β-Glu5* CRISPR/Cas9 edits (double mutants not characterized herein). Serendipitously, during the generation of *Dhr2* single edits, guide RNA placement produced in-frame edits (hereafter *ΔDhr2*; Figure 5A; methods) in five independent lines. All five T_0_ plants demonstrated a shared phenotype, showing extreme slow growth, stunted internode length (severe dwarfism), and unbranched female inflorescences, complete with husk leaves, in place of a tassel (Figure 5B). The T_0_ appearance of a consistent phenotype was unexpected and suggests that the *ΔDhr2* alleles act dominantly. How *ΔDhr2* acts dominantly is unknown, but diverse in-frame mutations have documented dominant effects (Chandrasekharappa *et al*., 1997; Tao *et al*., 2000; Reichenberger *et al*., 2001; Holland *et al*., 2007; Carmignac *et al*., 2012; Semler *et al*., 2012; Barthelson *et al*., 2021), often attributed to dominant-negative or gain-of-function alleles (Herskowitz, 1987; Veitia *et al*., 2018). These *ΔDhr2* edits strongly resemble *te1^-^* (Veit *et al*., 1998), albeit with more severe dwarfism and decreased vigour. All five *ΔDhr2* T_0_ plants exhibited very poor male and female fertility, together producing only 12 kernels. The vigour and viability of the T_1_ generation was worse, perhaps coincident with an increase in the edited allele dose; most *ΔDhr2* edited plants died as seedlings, and none produced offspring. This unexpected *te1^-^* phenocopy suggests that these dominant *ΔDhr2* edits cause the SAM to enter a developmental program akin to *te1^-^*.

### Extreme floral delay reveals genetic synergy between *id1^-^* and *te1^-^* mutants

Maize *te1^-^* loss of function alleles increase leaf number (Veit *et al*., 1998; Kawakatsu *et al*., 2006; Busche *et al*., 2023), yet the role of TE1 in flowering is uncharacterized. The phenocopy between *te1^-^* and our *ΔDhr2* edits suggests that these genes influence the same process. Due to the ties between *Id1* and *Dhr2* (Coneva *et al*., 2007; Minow *et al*., 2018; This report), we hypothesized that *Te1* and *Id1* may exist on parallel or linear genetic pathways. Therefore, to investigate the *Id1* and *Te1* genetic relationship, we crossed *id1^-^* and *te1^-^* single mutants and generated an F2 population (Figure 5C). If *Te1* acts downstream of *Id1* and *Dhr2* in a linear pathway, we expect *id1^-^ te1^-^* floral epistasis; specifically, the floral delay, or leaf number, of *id1^-^ te1^-^* should resemble that of the ‘high’ parent *id1*^-^. Indeed, *Dlf* was placed downstream of *Id1* by the fully epistatic floral delay observed in *id1^-^dlf ^-^* double mutants (Muszynski *et al*., 2006). Alternatively, if *Id1* and *Te1* influence leaf number through independent pathways, *id1^-^ te1^-^* leaf number would reflect the sum of the floral delays seen in the single mutants (additive inheritance). Exemplifying this, *PhytochromeB1*, *PhytochromeB2* and *Te1^-^* independently influence leaf initiation rate (plastochron) (Busche *et al*., 2023), with double mutants reflecting additive control of leaf initiation.

In the *id1*^-^ *te1*^-^ F2 population, we measured leaves produced before flowering as well as plastochron, as *Te1* impacts both total leaf number and the timing of leaf initiation (Veit *et al*., 1998; Busche *et al*., 2023). F2 plastochron was exclusively influenced by *Te1* genotype, with *Id1^+^ te1^-^* and *id1^-^ te1^-^* plants exhibiting an equally expedited plastochron and *id1^-^ Te1^+^* plants resembling *Id1^+^ Te1^+^* siblings (Figure 5D; Table S14). As previously reported(Veit *et al*., 1998; Busche *et al*., 2023)., *te1^-^* single mutants produced more leaves, and transition to spiral phyllotaxy near the floral transition (Figure 5CD leaf 18 in photo). *Id1^+^ te1^-^* exhibited tassel feminization, with shoots ending in the eponymous terminal ear. However, we observed varying penetrance, with only feminization of basal florets on some plants. Unexpectedly, we observed a non-additive synergistic (Pérez-Pérez *et al*., 2009) floral delay in *id1^-^ te1^-^* F2 plants, which produced a remarkable number of leaves (Figure 5D; Table S15; *id1^-^ te1^-^* make 90.00 ± 19.2 leaves, with 54.75 leaves predicted additively). Double mutants also produced aberrant apical inflorescences, with highly penetrant tassel feminization and husk leaves enveloping each apical inflorescence branch. Within these branching ‘terminal ears’ (Figure 5C), leafy shoots occasionally replaced kernels in ear-basal positions. Importantly, this increased leaf number exceeds that explained only by the production of terminal husk leaves; i.e., *id1*^-^ *te1*^-^ floral delay is mostly constituted of fully vegetative (non-husk) phytomers and cannot be explained by the homeotic changes to the apical inflorescence. Together, these results implicate *Te1* in autonomous maize flowering and, given *Te1* regulation of leaf primordia formation, could support a model where the cessation of leaf production influences maize flowering (Figure 5E; see discussion). Moreover, the genetic synergy observed in *id1*^-^ *te1*^-^ double mutants supports a non-linear genetic relationship between *Te1*, *Id1* and temperate maize floral induction. Specifically, the *id1*^-^ *te1*^-^ floral delay supports a genetic model where *Id1* and *Te1* control two independent pathways that converge on, or before, flowering.

## Discussion

Comparing the *id1^-^* and *Id1^+^* chromatin accessibility landscape revealed that nucleosome occupancy differences were enriched for TCP, MYB and AP2/ERF TF motifs, but not IDD/RID1 motifs. This suggests the remodelling of CRE chromatin accessibility occurs predominantly through *Id1* indirect effects on TCP, AP2/ERF and MYB TF signalling. While MYB TF motif enrichment is consistent with the documented protein-protein interaction between ID1 and MYB TFs (Zhou *et al*., 2026), how MYB TFs control nucleosome placement is poorly characterized. In contrast, the importance of TCP motifs in maize chromatin accessibility (Marand *et al*., 2025; Yao *et al*., 2026), and maize development (Doebley *et al*., 1997), is well documented. Additionally, yeast-2-hybrid analysis and *in vitro* pulldowns support orthologous interactions between RID1, OsTCP2, OsTCP10, and OsTCP11 (Zhang *et al*., 2022) and concurrent studies highlight maize ID1-TCP interactions (Zhou *et al*., 2026). Therefore, it seems plausible that ID1 may also alter TCP mediated nucleosome remodelling, some of which may control flowering or carbon metabolism.

Our motif enrichments also suggest *Id1* modulates chromatin accessibility via AP2/ERF TFs. AP2/ERF TFs are monocot-dicot conserved floral transition repressors (Chen, 2004; Jung *et al*., 2007; Mathieu *et al*., 2009; Yant *et al*., 2010; Lee *et al*., 2014; Zhang *et al*., 2022) (However, see Shim *et al*., 2022), with variation in domesticated maize flowering being controlled by the AP2/ERF floral repressor *ZmRap2.7* (Salvi *et al*., 2007). However, there is limited evidence of AP2/ERF TF chromatin accessibility modulation. Curiously, ChIP-seq demonstrated that Arabidopsis AP2 direct gene targets are equally partitioned between increased or decreased transcription in *ap2^-^* mutants (Yant *et al*., 2010). Nucleosome remodelling could explain this, with CREs that promote and repress transcription being contemporarily revealed by AP2 actions. Further, our results mirror those in rice, where RID1 directly regulates *OsERF136,* which represses *Hd3a* expression (Zhang *et al*., 2022). As such, AP2/ERF TFs acting downstream of *Id1* is an attractive hypothesis, and we support 68 AP2/ERF TFs as candidate direct ID1 targets. Since we highlight *AP2/ERF*-linked chromatin changes in the phloem, where florigenic PEBP signals are expressed (Chen *et al*., 2018), perhaps maize AP2/ERF TFs are likewise responsible for the *id1^-^* induced chromatin remodelling (Mascheretti *et al*., 2015) and repression of inductive PEBPs (Meng *et al*., 2011; Castelletti *et al*., 2020), precluding maize florigen production. However, AP2/ERF TF-family binding motifs sometimes exhibited elevated *Id1^+^* chromatin accessibility relative to *id1^-^* contexts, implying *Id1* mediated AP2/ERF TF network changes do not reflect simple repression of AP2/ERF signalling, as might be expected if this gene family strictly limits the maize floral transition.

Although IDD/RID1 motifs were not enriched within DARs, the RID1 motif was mildly enriched in DEG-linked ACRs, suggesting it may more accurately demarcate ID1 transcriptional targets than the longer IDD motif. Differences between *in vitro* determined TF motifs and *in vivo* chromatin targeting has been observed (Zhang *et al*., 2022; Bang *et al*., 2025), likely reflecting mechanisms (e.g. binding partners, post-translational modifications) that are difficult to replicate *in vitro,* or *in vitro* detection bias towards only high-affinity TF binding motifs (Khetan *et al*., 2025). Regardless, we observed stronger RID1 motif enrichments near DEGs, than DARs. We propose two hypotheses that could explain this: I) *id1^-^* secondary (indirect) effects alter chromatin accessibility more than transcription, better masking ID1 chromatin impacts or II) ID1 regulation strictly regulates RNA Polymerase II recruitment and it is exclusively downstream secondary effects that alter chromatin and CRE usage. Better characterization of ID1-interacting proteins may clarify how ID1 directly regulates transcription and/or nucleosome placement.

Filtering *id1^-^* DEGs for those with linked ACRs containing a RID1 TF motif, we provide a list of candidate direct ID1 targets. This DEG approach is an imperfect measure of ID1 targets, as it only encompasses expression differences at one time point in the day-night cycle. As the circadian clock affects maize flowering (Hung *et al*., 2012; Huang *et al*., 2017; Stephenson *et al*., 2019; Li *et al*., 2023; Zhao *et al*., 2023), and is perturbed in *id1^-^* mutants (Minow *et al*., 2018; This report), it is possible that the causative agents driving *Id1* flowering are missed by our approach. Future time course measurements would address this caveat, dovetailing well with our current analysis. Nonetheless, these candidate direct targets include floral regulator genes including *MADS67* (Zm00001eb327040) (Minow *et al*., 2018; Sun *et al*., 2020; Zhou *et al*., 2026), 24 *CCT* genes, and the *AP2/ERF* genes as discussed previously. Candidate ID1 targets also included genes that might explain the perturbed carbon metabolism seen in *id1^-^* plants. These included *XylA* (Zm00001eb265380), which has elevated expression in *id1^-^* plants. *XylA* interconverts D-Glucose to D-Fructose (Stevis and Ho, 1985) which may explain the elevated fructose in *id1^-^* developing leaves (Coneva *et al*., 2012). A non-exclusive alternative hypothesis is that, since *XylA* also catalyzes the interconversion of D-Xylose to D-Xylulose (Stevis and Ho, 1985), it may feed excess xylose to glycolysis via xylulose kinase (Zm00001eb094750) and the pentose-phosphate pathway. We also find putative ID1 binding sites near a family of *β-Glu* genes that are known to dramatically lose expression in *id1^-^* (Coneva *et al*., 2007; Minow *et al*., 2018; This report). Although the maize substrate targets of these β-GLU remain uncharacterized, they are closely related to *Sorghum bicolor Dhurrinase* (*SbDhr*) genes. *SbDhr* acts on dhurrin, a highly abundant (up to 30% of dry shoot biomass) (Halkier and Møller, 1989) cyanogenic glucoside that deters herbivory. Intriguingly, dhurrin can be cycled to provide carbon and nitrogen skeletons via an emerging *S. bicolor* pathway that requires GST-mediated glutathione conjugation to dhurrin, preventing cytotoxic cyanogenesis (Bjarnholt *et al*., 2018; Gleadow *et al*., 2021). Given that our multimodal genomics reveals extensive alterations in glutathione metabolism genes, it may be that a yet unknown substrate of the maize β-GLUs is likewise cycled to move carbon and nitrogen within the developing leaf.

Through in-frame CRISPR/Cas9 editing of one of these uncharacterized *β-Glu* genes, *Dhr2*, we unexpectedly observed a dominant *te1^-^* phenocopy. This *ΔDhr2* phenotype precipitated reexamination of the role of *Te1* in the reproductive transition, prompting an *id1^-^* epistasis check to see if SAM-expressed *Te1* could be involved in *Id1* signalling. Double mutant *id1^-^ te1^-^* plants exhibited a synergistic floral delay, composed of fully vegetative (i.e. non-husk) phytomers, highlighting a role for *Te1* in controlling maize flowering on a genetic pathway independent of, but convergent with, *Id1*. Loss of function *te1^-^* alleles have been previously reported to increase the number of leaves produced before flowering (Veit *et al*., 1998; Busche *et al*., 2023). However, several factors concealed this *bona fide* floral delay. Firstly, *te1^-^* homeotic changes induce apical husk leaf formation masking the underlying floral delay. This mirrors Arabidopsis *ap2* mutants, whose strong homeotic defects concealed its roles in the floral transition for many years (Chen, 2004; Yant *et al*., 2010). Additionally, the *te1^-^* shortened plastochron causes a similar days-to-anthesis timing. However, days-to-anthesis, or days-to-flowering, is a composite trait that pairs a genetically determined developmental plan with a more environmentally influenced, or plastic, growth rate (Wallace, 1985; Koornneef *et al*., 1991). Our *te1^-^* work reinforces that leaf number is a robust measure of developmental flowering time.

The floral homeotic consequences of *te1* mutations (Veit *et al*., 1998; Sun *et al*., 2024) mirror that seen in mutants that control both reproductive identity and timing (Mandel and Yanofsky, 1995; Chen, 2004; Liu *et al*., 2009; Yant *et al*., 2010), perhaps tying TE1 repression of leaf formation to reproductive meristem formation and flowering time. Normally, during the maize floral transition, leaf formation stops, the SAM enlarges and elongates, and reproductive meristems emerge (Abbe *et al*., 1951; Irish and Nelson, 1991; Meng *et al*., 2011). We speculate that *te1^-^* continual leaf formation inhibits SAM enlargement and/or the organizational changes needed to produce reproductive structures. Specifically, as plant cell fate is heavily influenced by positional effects (Scheres, 2001; Pallakies and Simon, 2010), persistent recruitment of peripheral SAM cells into leaf fates may constrain the undifferentiated stem cell pool size, precluding the establishment of the *ZmBIF3*/*ZmCLE7* (orthologs of *WUS*/*CLV3*) stem cell niches needed for reproductive meristem formation (Rodriguez-Leal *et al*., 2019; Chen *et al*., 2021; Liu *et al*., 2021; Bang *et al*., 2025; Xu *et al*., 2025). Supporting this model, larger SAM size has been correlated with earlier maize flowering (Leiboff *et al*., 2015; Thompson *et al*., 2015). This SAM competition between leaf and reproductive meristem cell identities mirrors current understanding of inflorescence development, where models stress intra-SAM competition for founder cells and/or growth metabolites between bracts (modified leaf primordia) and nascent reproductive meristems (Coen and Nugent, 1994; Baum and Day, 2004; Chuck and Bortiri, 2010; Whipple *et al*., 2010; Xiao *et al*., 2022). Similarly, perhaps the inability to pause vegetative-phase leaf primordia initiation precludes the formation of reproductive meristems.

Genetic synergy, like that observed here between *id1^-^ te1^-^*, typically indicates that two loci converge on the same pathway (Pérez-Pérez *et al*., 2009) and compensate for one-another in single mutant states. The pitfall with genetic synergy is misinterpreting an additive genetic relationship (Pérez-Pérez *et al*., 2009). Here, however, pronounced deviation from the expected additive floral delays strongly supports genetic synergy between *Te1* and *Id1* pathways. This produces two potential models where *Te1* and *Id1* converge on flowering, or leaf formation (Figure 5E). We favour convergence at leaf formation because: I) *ΔDhr2* edits phenocopied *te1^-^*, implying altered leaf initiation as in *te1^-^* II) it is a more parsimonious model, involving fewer interactions and III) it may reconcile why only part of the *id1^-^* floral delay appears to act through *Zcn*-*Dlf1* induction (Muszynski *et al*., 2006; Meng *et al*., 2011; Zhou *et al*., 2026), as perhaps *Id1* both promotes floral inductive signals and inhibits leaf formation through different pathways. If *Id1* helps repress leaf formation, it may exacerbate the *te1^-^* impacts on leaf primordia formation, and flowering, as described above. How maize immature leaves may regulate leaf initiation is unclear, but immature rice leaves express *PLASTOCHRON1* (Itoh *et al*., 1998), which non-autonomously inhibits leaf formation and affects rice SAM size (Itoh *et al*., 1998; Miyoshi *et al*., 2004). Mirroring the maize independence of *Id1* and *Te1* pathways, rice *PLASTOCHRON1* controls leaf initiation independently of the rice TE1 ortholog, *PLASTOCHRON2* (Kawakatsu *et al*., 2006). *PLASTOCHRON1* encodes a cytochrome p450 enzyme (Miyoshi *et al*., 2004) with unknown substrate specificity. Cytochrome p450s act on diverse substrates including cyanogenic glucosides, like dhurrin (Laursen *et al*., 2015). A maize *PLASTOCHRON1* ortholog similarly affects leaf growth (Sun *et al*., 2017), but its mechanisms remain mysterious (Laureyns *et al*., 2022). Taken together, the extreme floral delay of *id1^-^ te1^-^* plants highlights *Te1* autonomous flowering roles, and *Id1-Te1* crosstalk, while hinting at a potential connection between the cessation of leaf formation and reproduction.

## Methods

### Controlled plant growth, tissues and nuclei isolation and library generation

Siblings segregating the *id1-m1* allele (Colasanti *et al*., 1998) introgressed into the B73 background were growth-chamber-grown (Conviron) under a 16-hour photoperiod at a light intensity of ∼1000 μmolm^2^s^-1^, with daytime temperatures of 28°C and night temperatures of 22°C. Siblings were genotyped using DNA extraction and PCR conditions as previously described (Wong and Colasanti, 2007). To ensure accurate genotypes, all plants were genotyped twice from independent DNA extractions, and only samples with concordant genotypes were used for downstream analysis. At one hour post dawn, V7 stage 2cm long immature leaf cylinders were harvested from 3cm above the last visible maize node. From half this cylinder, nuclei were extracted for two biological scATAC-seq replicates from plants homozygous for either the *id1-m1* (*id1^-^*) or *Id1-B73* (*Id1^+^*) allele. Nuclei for scATAC-seq were extracted through modification of established methods (Sikorskaite *et al*., 2013). First the fresh immature leaf tissue was chopped vigorously for 2 minutes in nuclear extraction buffer (1X NIB: 10mM MES:KOH pH 5.4, 10mM NaCl, 10mM KCl, 2.5mM EDTA pH 8, 250 mM sucrose, 0.1mM spermine, 0.5mM spermidine, 1mM fresh DTT, 1% vol/vol BSA) with 0.5% vol/vol Triton X100. This chopped tissue was passed through a 40μm cell strainer (pluriselect) into a 2mL conical tube and spun at 4°C and 500g for 5 minutes. The supernatant was removed, and the nuclei pellet was then resuspended in 500μL 1X NIB. The resuspended nuclei then pass through a 10 μm cell strainer (Pluriselect) before being gently layered on top of 1mL of 35% vol/vol Percoll:1XNIB in another 2mL conical tube and spun for 10 minutes at 4°C and 500g. All supernatant was again removed, and the nuclei pellet resuspended in 1X TAPS buffer (10mM TAPS-NaOH, pH8.0, 5mM MgCl_2_) and stained with DAPI (1μg/mL final concentration) before being counted in a hemocytometer. 15,000 nuclei were then Tn5 tagmented per replicate as previously described (Ricci *et al*., 2019), except using 1.75μg Tn5. These tagmented nuclei were then used to generate 10X scATAC-seq libraries (10X Genomics Chromium Next GEM Single Cell ATAC Reagent Kits v1.1), in place of their proprietary Tn5 tagmentation reagents, but otherwise following manufacturer protocols. The ensuing libraries were sequenced (∼400 million 150 bp paired end reads) on an Illumina NovaSeq 6000.

For snRNA-seq nuclei isolation and purification were conducted according to a previously described method with minor modification (Zhang *et al*., 2024b). Briefly, frozen immature leaf samples were directly chopped on ice for approximately 1 minute using 600μL of pre-chilled 1X NIB containing 0.4U/μL RNase inhibitor (Roche, Protector RNase Inhibitor, Cat. RNAINH-RO) and a 0.1% NP-40. Following chopping, the entire mixture was passed through a 40μm cell strainer (Pluriselect) and centrifuged at 4°C and 500g for 10 minutes. The supernatant was carefully decanted before the nuclei pellet was resuspended in 500μL of a wash buffer, comprising 10mM MES-KOH (pH 5.4), 10mM NaCl, 250mM sucrose, 0.5% BSA, and 0.2U/μL Roche RNase inhibitor. The nuclei were then filtered through a 20μm cell strainer and gently layered onto the surface of 1 mL of a 35% Percoll 65% NIB buffer in a 1.5-mL centrifuge tube before centrifugation at 4°C and 500g for 10 minutes. Then the supernatant was removed, and the pellets were washed once in 650μL of Nuclei Wash C Resuspension buffer (NWR buffer: 1% BSA, 0.2U/μL RNase inhibitor, 1xPBS) before being pelleted at 4°C and 500g for 10 minutes and final resuspension in 50 μL NWR buffer. Approximately 5μL of nuclei were diluted tenfold and stained with DAPI (Sigma Cat. D9542) to evaluate nuclei quality and quantity under a fluorescence microscope. These nuclei were diluted with DNB buffer to a final concentration of 1,000 nuclei/μL, with a total of ∼16,000 nuclei used as input for 10X Chromium Next GEM Single Cell 3’GEM Kit v3.1 (Cat# PN-1000123) library generation, following the manufacturer’s instructions (10xGenomics, CG000315_ChromiumNextGEMSingleCell3-_GeneExpression_v3.1_DualIndex_RevB). The libraries were subsequently sequenced using the Illumina NovaSeq 6000 in dual-index mode (∼200 million 150 bp paired end reads).

### scATAC-seq processing

Raw BCL files were first demultiplexed into FASTQ format using the 10x Genomics tool cellranger-atac makefastq (v2.1.0). Subsequent preprocessing was performed with *cellranger-atac count.* During this step, cell barcodes were corrected against a 10x barcode whitelist. Paired-end reads from the four samples (MU5, MU9, WT2, and WT7) were trimmed, filtered using a MAPQ score of greater than 30 and aligned to B73 v5 of *Zea mays* reference genome using Cell Ranger ATAC’s algorithm. The resulting position sorted BAM files were used for downstream analysis (Hufford *et al*., 2021). Mitochondria and chloroplasts assemblies were included in the reference genome for quality control and later exclusion. Paired-end, uniquely mapped reads were filtered via SAMtools (Li *et al*., 2009) to retain quality of greater than or equal to 10, before deduplication using Picardtools, leaving unique Tn5 insertion sites for each replicate. Organellar reads were removed and Tn5 insertions further filtered against a maize blacklist (Supplemental Data S2), which outlines sequences of the genome with enzymatic Tn5 bias, which would cause false-positive ACR detection if not removed. Aligned BAM files were converted to single base pair resolution Tn5 integration sites in BED format by shifting read coordinates by +4 bp on the positive strand and -5 bp on the negative strand, retaining only unique insertion events per barcode. Socrates (Marand *et al*., 2021) was used to identify high-quality barcodes by generating log10-transformed barcode rank plots of Tn5 integration counts per barcode, with thresholds defined at the knee inflection point for each sample. More stringent quality control thresholds than default were applied, including increased TSS enrichment (z ≥ 3), higher minimum cutoffs for TSS enrichment and FRiP, and a cap on the total number of cells, to improve signal-to-noise and retain high-confidence nuclei.

### ACR identification

ACRs were identified from single-base resolution Tn5 insertion BED files. For each sample, insertion sites were randomly partitioned into two pseudo-replicates of equal size. MACS2 (Zhang *et al*., 2008) peak calling was performed on both pseudo-replicates and the full dataset using the parameters -g 1.6e9 --extsize 75 --shift -50 --nomodel --keep-dup all --qvalue 0.01. Reproducible ACRs were defined as peaks detected in the full dataset that overlapped peaks from both pseudo-replicates by ≥10% of their length. Peak summits corresponding to reproducible ACRs were expanded by ±250 bp to generate fixed-width regions of 500bp summits. ACRs from all samples were then merged and overlapping regions were filtered to generate a master peak set.

### Clustering and cell type annotation

Clustering was performed using ArchR (Granja *et al*., 2021) to group chromatin accessibility profiles based on similarity. Dimensionality reduction was carried out using the *addIterativeLSI* function with a low clustering resolution (0.2) to capture broad cell populations for comparison between *Id1^+^* and *id1^-^* samples. Cells were then clustered using the Louvain algorithm with a correlation cutoff of 0.6 and bias correction was applied using TSS enrichment, log10-transformed fragment counts, FRiP, and doublet enrichment scores (Figure S1). A Uniform Manifold Approximation and Projection (UMAP) (Mcinnes *et al*., 2018) embedding was generated for visualization of the clustered cells.

Cluster identities were annotated based on a predefined set of marker genes curated from the literature (Figure S3). Marker gene accessibility was visualized using dot plots generated from ArchR (Granja *et al*., 2021) gene score matrix values, where scaled average accessibility and the percentage of cells exhibiting accessibility were calculated for each gene. Differential chromatin accessibility analysis was performed on the peak count matrix generated using featureCounts from the master peak set above. Two pseudoreplicates per cluster were generated by randomly sampling two-thirds of reads without replacement, with peaks within the introgression region (chr1:236,615,096–252,058,603) excluded from analysis. Differential chromatin accessibility between *Id1^+^* and *id1^-^* samples was assessed both globally and within individual cell clusters using DESeq2 (Love *et al*., 2014). Peaks with an adjusted p-value (FDR) < 0.05 were considered significantly differentially accessible and results were reported as *Id1^+^* vs *id1^-^*. Differentially accessible regions were subsequently subjected to functional enrichment analysis using FGSEA in R. Gene ontology (GO) terms were considered significantly enriched at FDR < 0.05. Cell type was assigned based upon a marker gene expression (snRNA-seq) and gene-body accessibility (scATAC-seq) combined with gene ontology enrichment comparison of expression or gene-body accessibility of each cell-type cluster to the rest of the cells. As *Id1^+^* and *id1^-^* cell clusters were largely unchanged (Figure S2), all annotation was conducted on *Id1^+^* tissue, as *id1^-^* perturbed vasculature gene signatures (Figure S4). Further, unlike previous maize leaf profiles (Marand *et al*., 2021; Yan *et al*., 2025a), the photosynthesis-coupled signature that delimitates mature bundlesheath cells was absent, likely due to their immature state. Instead, developing bundlesheath identity was disentangled from mesophyll cells by relative *Shortroot* (Zm00001eb020850; Zm00001eb326020) gene-body accessibility/expression enrichment (Chang *et al*., 2012; Xu *et al*., 2021). Additionally, the cell-type midrib parenchyma was determined strictly by process of elimination, as it exhibited weak gene ontology signatures and contains no known cell-type specific genes. Similarly, a paucity of cell-type specific characterization complicates our annotation of certain cell types; specifically, those arising from phloem sub-functionalization, vasculature parenchyma and some protoderm stages were somewhat ambiguous, and this uncertainty has left us with vague annotations including the scATAC-seq clusters “intermediate maturity protoderm1/2”. Future characterization may improve or alter these specific cell-type calls.

### Motif discovery and enrichment

Position weight matrices (PWMs) were obtained from the JASPAR 2024 database (Rauluseviciute *et al*., 2024). Due to the limited availability of curated motif datasets for maize, position weight matrices (PWMs) derived from Arabidopsis were used to identify homologous motifs in the *Zea mays* genome for ID1, ERF, and TCP, based on the family-level conservation of transcription factor binding (Rogers and Bulyk, 2018; Baumgart *et al*., 2025). Motif occurrences were identified genome-wide using FIMO (Grant *et al*., 2011) with --max-stored-scores set to 500,000 and a threshold of 0.05, with the exception of the RID1 motif (Zhang *et al*., 2022), for which --max-stored-scores was increased to 1,000,000. Each motif list was filtered independently to remove low-confidence matches (where p-values plateaued) and to exclude ACRs with fewer than two Tn5 insertion sites. The resulting motif coordinates were intersected with ACRs using BEDTools (Quinlan and Hall, 2010) to identify motif-containing regions and assess their proximity to differential ACRs and nearby genes. *De novo* motif discovery was additionally performed using STREME (Bailey, 2021) with the following parameter. Motif accessibility was quantified using ChromVAR (Schep *et al*., 2017) which computes GC bias-corrected deviation scores for each motif across single cell. These scores were then projected onto the UMAP to visualize motif accessibility score across cell types.

### snRNA processing and Clustering

Raw FASTQ files were processed using Cell Ranger (v8.0.0). The *Zea mays* B73 reference genome (v5.0) was used for alignment, and sequencing reads from four samples (MU5, MU9, WT2, and WT7) were processed using the *cellranger count*. Barcode filtering was further refined using Socrates (Marand *et al*., 2021) to retain high-quality nuclei, applying criteria consistent with those described for scATAC-seq preprocessing. During tandem Seurat (Hao *et al*., 2021) object generation, sample-specific minimum feature thresholds were defined based on barcode rank inflection points and set to 260 (MU5), 330 (MU9), 430 (WT2), and 260 (WT7). Following normalization and identification of 2,000 variable features, the data were scaled and subjected to principal component analysis (PCA). A shared nearest neighbor graph was constructed using the first 15 principal components, as determined by inspection of the elbow plot, and clustering was performed at a resolution of 0.5. QC indicated that cluster 1 predominantly represented low-quality or technical artifacts and was therefore excluded from downstream analyses (Figure S5). Cluster 9 contained a mixture of xylem and phloem vasculature parenchyma cell types and was subsequently subjected to sub-clustering to resolve this heterogeneity.

### snRNA cell type annotation

Gene expression patterns across clusters, both globally and within genotype specific subsets, were visualized using Seurat’s (Hao *et al*., 2021) DotPlots of marker gene expression were generated with the marker genes as described above, representing both average expression and the proportion of expressing cells (Figure S6). Raw count matrices from each sample-specific RNA layer were extracted and combined with cluster assignments and sample identity. Counts were aggregated by summing expression values across cells within each gene, cluster, and sample combination to generate pseudobulk profiles. Pseudobulk replicates were generated as described for scATAC-seq, with three replicates constructed per sample cluster combination, followed by quantile-quantile normalization. The resulting matrices were combined to produce a final gene by cluster sample count matrix for downstream differential analysis. Both global and cluster-specific differential analyses were performed using a false discovery rate (FDR) threshold of 0.05 and the introgression region removed.

### CRISPR/CasG editing and *te1^-^ id1^-^* double mutant analysis

Stable maize transformation was performed on immature embryos as previously published, but with minor modifications (Masters *et al*., 2020; Chen *et al*., 2026). Briefly, two gRNAs targeting the *Dhr2* (Zm00001eb138340) coding sequence were cloned into a PGGB vector, via HiFi Assembly (NEB). PGGBFigurederived from the pBEU11 backbone but includes two morphogenic regulator genes: the maize transcriptional regulator BABY BOOM (ZmBBM/EREB53) and a wheat GROWTH REGULATING FACTOR 4–GRF-INTERACTING FACTOR 1 (GRF4:GIF1) fusion protein, which together enhance maize transformation efficiency and regeneration (Chen *et al*., 2026). PGGB also expresses a *CASS* cassette under the Maize *Ubiquitin1* promoter and two gRNAs under the *OsU3* and *OsUCP2* promoter respectively (Chen *et al*., 2026) to induce deletion edits. Complete binary constructs were introduced into *Agrobacterium tumefaciens* strain EHA105 via electroporation. To generate the in-frame *ΔDhr2* deletion, two promoters, *OsU3P* and *OsU6P2*, were used, with each promoter driving the expression of one gRNA. The two guide sequences were CAGCCGCCATGATAATGGC and TGGTATGAAGCTAAGGCTA. Using these guides, we recovered five 66-bp in-frame deletions near the beginning of the first exon, starting at the 70th nucleotide of the coding region. Double edits of the linked *Dhr2* and *β-Glu5* genes are not characterized as of writing.

For immature embryo collection, B104 inbred plants were grown under greenhouse conditions before embryo harvest 10–14 days after pollination. These immature embryos were immediately transformed with the EHA105 Agrobacterium carrying the final binary vector. Following Agrobacterium co-cultivation, transgenesis selection via 0.3% glufosinate-ammonium (basta; Liberty brand), shoot induction and root induction, regenerated plantlets were transferred to a Percival growth chamber for acclimatization for up to two weeks before greenhouse transfer for growth until seed production. Transgene integration and edit characterization were confirmed by PCR using construct-specific primers (Table S16).

To investigate the genetic interactions between *Id1* and *Te1*, greenhouse grown *id1-m1* (*id1^-^*) B73 introgressed plants (Colasanti *et al*., 1998; Minow *et al*., 2018) were hand-crossed with *te1-1* (*te1^-^*) (Matthews *et al*., 1974; Veit *et al*., 1998) plants from an out-bred background as provided by the Maize Genetics Cooperative Seed Stock Center (supported by USDA-ARS). F2 seed collected from a single F1 family were grown to compare single and double mutants to their *Id1^+^* and/or *te1^+^* siblings. Total leaf number and leaf initiation rate (determined as the rate of collared leaf emergence up until tassel emergence) were quantified in two independent greenhouse grow outs of the same F2 population in 2025 and 2026. Mutant *id1^-^* plants were determined via established PCR genotyping (Wong and Colasanti, 2007) while *te1^-^* plants were established based upon apical inflorescence feminization (terminal ear formation), as no *te1-1* molecular genotyping assay exists. In the first grow out, *id1^-^ te1^-^* double mutants were observed at the expected 1/16^th^ rate (4/52; Chi squared test; p=0.6674). To bolster our statistical power, we later increased the number of F2 *te1^-^* plants by selecting for young seedlings with elevated plastochron in the second grow out. This seedling selection elevated the prevalence of *te1^-^ id1^-^* double mutants but distorted the segregation ratios. Between these two grow outs, 92 individuals were grown to flowering and phenotyped. No major changes were observed between grow outs.

## Supplement list

Figure S1 – scATAC-seq (FRiT), fraction of reads in peaks (FRiP), organellar reads, and doublet scores.

Figure S2 – scATAC-seq clusters had an equal number of *Id1^+^* and *id1^-^* cells

Figure S3 – scATAC-seq cell-type markers dot plots

Figure S4 – scATAC-seq of *id1^-^* cells and the corresponding cell-type enrichments of *Sucrose Transporter* (*SUT)* gene-body chromatin accessibility

Figure S5 – snRNA-seq median UMI and unique gene counts

Figure S6 – snRNA-seq cell-type markers dot plots

Figure S7 – Chromatin accessibility of IDD-family TF motifs in immature leaf cell types

Figure S8 – *Id1^+^* vs *id1^-^* transcription factor binding motif chromatin accessibility changes

Table S1 – Gene ontology (GO) enrichments within scATAC-seq clusters

Table S2 – *Id1^+^* vs *id1^-^* comparison of gene-body chromatin accessibility over the Indeterminate1 locus

Table S3 – *Id1^+^* vs *id1^-^* differentially accessible regions (DARs) in each cell-type

Table S4 – Fast gene set enrichment (FGSEA) for DAR-linked genes

Table S5 – Genes linked to DARs

Table S6 – Floral regulator genes linked to DARs

Table S7 – *Id1^+^* vs *id1^-^* differentially expressed genes (DEGs) in each cell-type

Table S8 – Floral regulator differentially expressed genes (DEGs)

Table S9 – Cell-type enrichments of differentially expressed floral regulator genes

Table S10 – *Id1^+^* vs *id1^-^* DAR motif enrichment compared to the global ACR motif complement.

Table S11 – Fast gene set enrichment (FGSEA) for DAR-linked genes with AP2/ERF and TCP motifs

Table S12 – *Id1^+^* vs *id1^-^* DEG-linked ACR motif enrichment compared to the global ACR motif complement.

Table S13 – Candidate ID1 target genes.

Table S14 – F2 id1-te1-leaves per day (plastochron)

Table S15 – F2 id1-te1-total leaves produced before flowering

Table S16 – ΔDhr2 genotyping primer sequences

Data S1 – All IDD/RID1 motif calls

Data S2 – V5 Maize genome Tn5 blacklist

## Acknowledgements

This research was supported by the National Science Foundation (IOS-2026554 and IOS-1856627) to RJS as well as the UGA Office of Research to RJS. A.P.M. was supported by the National Institute of General Medical Sciences of the National Institutes of Health (1R00GM144742). We would like to thank Micheal Boyd and Noah Herrington for growth facility maintenance and unparalleled plant care, as well as the staff at the Georgia Advanced Computing Resource Center (GACRC) and Compute Canada for their excellent computing clusters. We also thank our talented lab technicians Yinxin Dong and Yangyang Xu for their hard work that keep the schmitz lab running smoothly.

